# A Golden Gate-based Plasmid Library for the Rapid Assembly of Biotin Ligase Constructs for Proximity Labelling

**DOI:** 10.1101/2021.10.15.464533

**Authors:** Kevin Goslin, Andrea Finocchio, Frank Wellmer

## Abstract

Proximity-labelling has emerged as a powerful tool for the detection of weak and transient interactions between proteins as well as the characterization of subcellular proteomes. One proximity labelling approach makes use of a promiscuous bacterial biotin ligase, termed BioID. Expression of BioID (or of its derivates TurboID and MiniTurbo) fused to a bait protein results in the biotinylation of proximal proteins. These biotinylated proteins can then be isolated by affinity purification using streptavidin-coated beads and identified by mass spectrometry. To facilitate the use of proximity-labelling in plants, we have generated a collection of constructs that can be used for the rapid cloning of TurboID and MiniTurbo fusion proteins using the Golden Gate cloning method. To allow for the use of the constructs in a range of experiments we have designed assembly modules that encode the biotin ligases fused to different linkers as well as different commonly used subcellular localization sequences. We demonstrate the functionality of these vectors through biotinylation assays in tobacco (*Nicotiana benthamiana*) plants.

## INTRODUCTION

Cellular processes are executed and controlled through interacting proteins. The characterization of protein-protein interactions (PPIs) is therefore a critical step in the understanding of biology. Conventional approaches such as affinity purification followed by mass spectrometry (AP-MS) or yeast-2-hybrid (Y2H) face a number of limitations including the isolation of interaction partners under non-physiological conditions following cell lysis or a dependency on high affinity interactions. In recent years, proximity-labelling has emerged as a powerful tool that allows for the detection of weak and transient interactions between proteins as well as the characterization of subcellular proteomes (reviewed in [1, 2]).

One proximity labelling tool makes use of a promiscuous biotin ligase enzyme, termed BioID, which carries a point mutation in the biotin ligase BirA from *Escherichia coli* [3]. Expression of BioID fused to a bait protein in cells results in the biotinylation of proximal proteins which are either direct interactors or nearby proteins that do not directly interact with the fusion protein. As biotinylation is a rare occurrence in most organisms these biotinylated proteins can then be isolated by affinity purification using streptavidin coated beads and identified by mass spectrometry (MS) (Fig. 1A). Exogenous biotin is typically required for labelling of proteins so that labelling time is controlled by the addition of biotin to the cell, with labelling time generally between 15 – 18 h [3, 4]. Recently, novel variants of BioID have been developed through a directed evolution approach [5]. One of these mutant proteins, termed TurboID, displays a much faster rate of biotinylation in cells with labelling occurring within 10 min of biotin addition [5]. TurboID displays a higher level of activity at lower temperatures than BioID, possibly because BioID was derived from *E. coli* while TurboID was evolved in yeast [5]. In the same study researchers developed another highly active variant of this biotin ligase, termed MiniTurbo, which lacks the N-terminal (Nt) domain of TurboID resulting in a smaller variant (28 versus 35 kDa). MiniTurbo displays reduced stability [4, 5] but may produce a lower level of background biotinylation in the absence of exogenous biotin compared to TurboID [5]. The properties of these variants therefore allow for the capture of highly dynamic protein interactions at lower temperatures. To date TurboID has been shown to be active in a number of organisms and at a range of temperatures, including in *Drosophila melanogaster, Caenorhabditis elegans, Arabidopsis thaliana, Nicotiana benthamiana* and mammalian cells [5-8].

**Fig 1.**
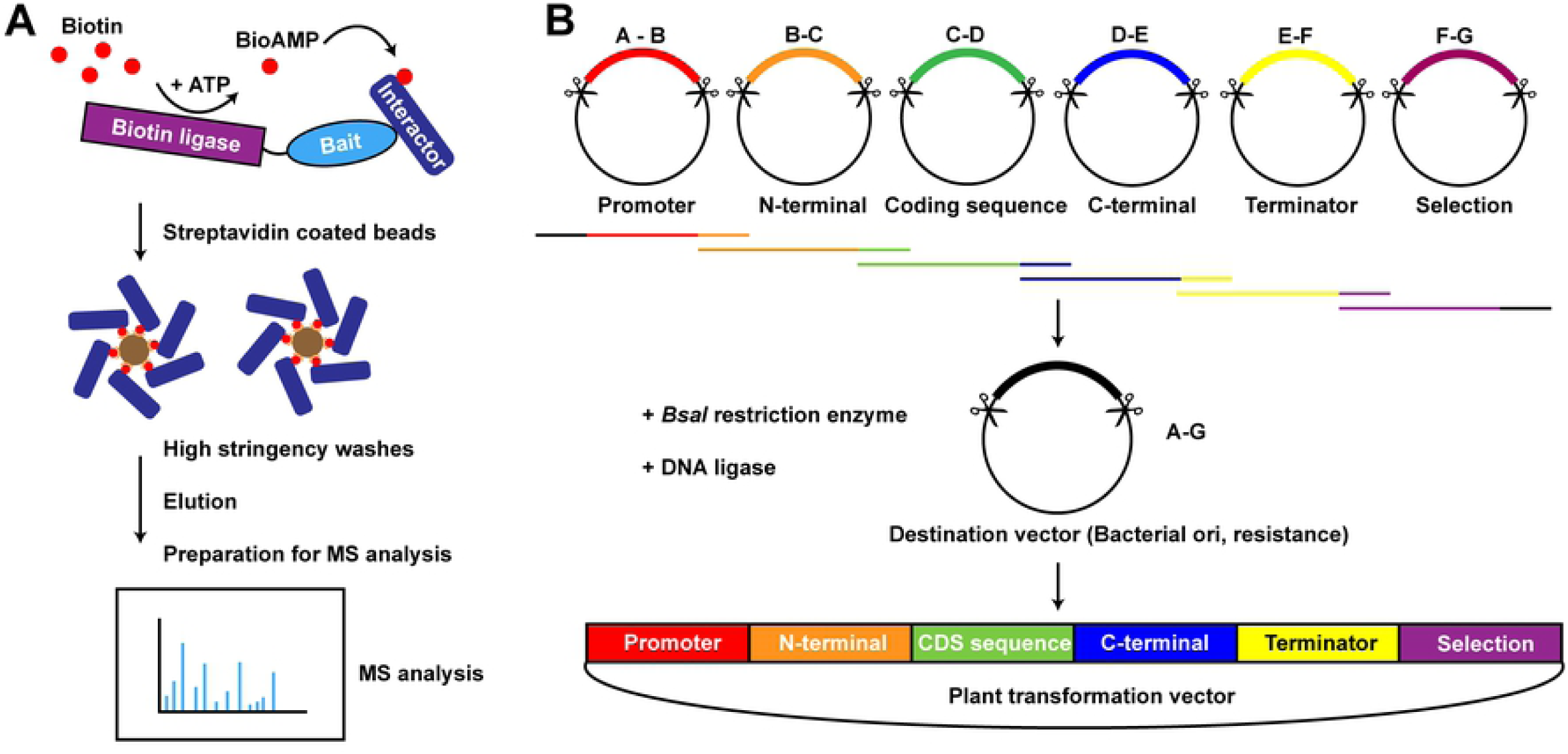
Overview of biotin ligase proximity labelling and GreenGate cloning. **(A)** Activity of a promiscuous biotin ligase results in the generation and release of reactive bioAMP and the biotinylation of proximal proteins. Biotinylated proteins can be isolated by affinity purification using streptavidin coated beads. High stringency washes are performed to remove background. Following elution of proteins from the beads peptides can be identified by mass spectrometry. **(B)** The Green Gate system of cloning utilizes a Golden Gate assembly approach. Entry vectors are first generated that contain single elements flanked by *Bsa*I recognition sites and one of seven different overhangs (A-G). Upon digestion with *Bsa*I individual elements are released from the entry vectors. Annealing of complementary overhangs and T4 ligase catalyzed DNA ligation results in the assembly of a final expression construct. Panel adapted from [16].

Over the last five decades a combination of type-II restriction enzymes and DNA ligases has been employed to generate recombinant plasmids. Although effective, this method can be time-consuming and may require multiple rounds of cloning steps. To overcome this limitation a number of cloning strategies have been developed in order to facilitate the rapid cloning of constructs. These include approaches making use of recombination-based cloning methods such as the Univector plasmid-fusion system, based on the Cre-*lox* system [9] or the widely used Gateway technology that makes use of the lambda phage DNA integration machinery [10, 11]. Although the Gateway system allows for the efficient insertion of multiple genes in to a single vector and many Gateway compatible vectors now exist, there are a number of disadvantages to the system including the high cost of the enzymes required for the reactions and the generation of a 25 bp ‘scar’ that is left flanking the integrated DNA as a result of the recombination site sequence. Another popular cloning strategy, termed the Golden Gate method, does not rely on recombination but instead makes use of type-IIS restriction enzymes [12]. These enzymes possess a specific DNA recognition site but cleave the DNA outside of this site, allowing for the generation of restriction site overhangs that can contain a customized sequence of nucleotides. Individual cloning modules are first generated as sequences flanked by two type-IIS restriction enzyme sites containing complementary overhangs with the desired preceding and succeeding modules. Upon digestion with the single type-IIS restriction enzyme the modules yield complementary overhangs allowing for their ligation in the presence of DNA ligase (Fig. 1B). With this method different combinations of individual modules can be quickly assembled together in a single reaction. Golden Gate cloning has been employed in a large number of studies and several resources exist including cloning kits such as the GoldenBraid [13, 14] and Golden GateWay series of vectors [15]. One cloning kit, termed GreenGate, was designed to facilitate rapid cloning of constructs using commonly used elements of plant transformation vectors [16]. Assembly modules are made up of six different types of element which typically include a promoter, N-terminal tag, coding sequence, C-terminal tag, terminator and resistance sequence. Cleavage of these modules with the type-IIS restriction enzyme yields fragments with overhangs designated type A-G which are then ligated in to a backbone destination vector containing bacterial resistance and overhang type A-H, forming a higher order assembly of plasmid [16].

Here, we present a collection of constructs that have been designed for the rapid cloning of the TurboID and MiniTurbo biotin ligases using the Golden Gate cloning method which are compatible with the GreenGate series of vectors. To facilitate a range of experiments we have designed assembly modules that encode the biotin ligases fused to different linker-lengths as well as different commonly used localization sequences. To demonstrate the functionality of these vectors we have used elements of the collection to generate plant transformation vectors and transiently expressed them in tobacco plants (*Nicotiana benthamiana*). We demonstrate that the biotin-ligase fusion proteins encoded by these vectors localize to the expected subcellular compartment and can biotinylate proximal proteins *in vivo*.

## RESULTS AND DISCUSSION

### Generation of a biotin ligase cloning library

In order to facilitate the rapid cloning of biotin ligase constructs we generated a library of 26 entry level cloning modules (Table 1). Each module is comprised of a TurboID or MiniTurbo coding sequence flanked by *Bsa*I restriction enzyme sites. Modules were designed to allow for the enzyme to be positioned at the beginning, middle or end of a construct (overhangs B-C / C-D/ D-E, respectively) following a promoter sequence, as per the Green Gate cloning kit layout [16] (see Materials and Methods).

**Table 1:**
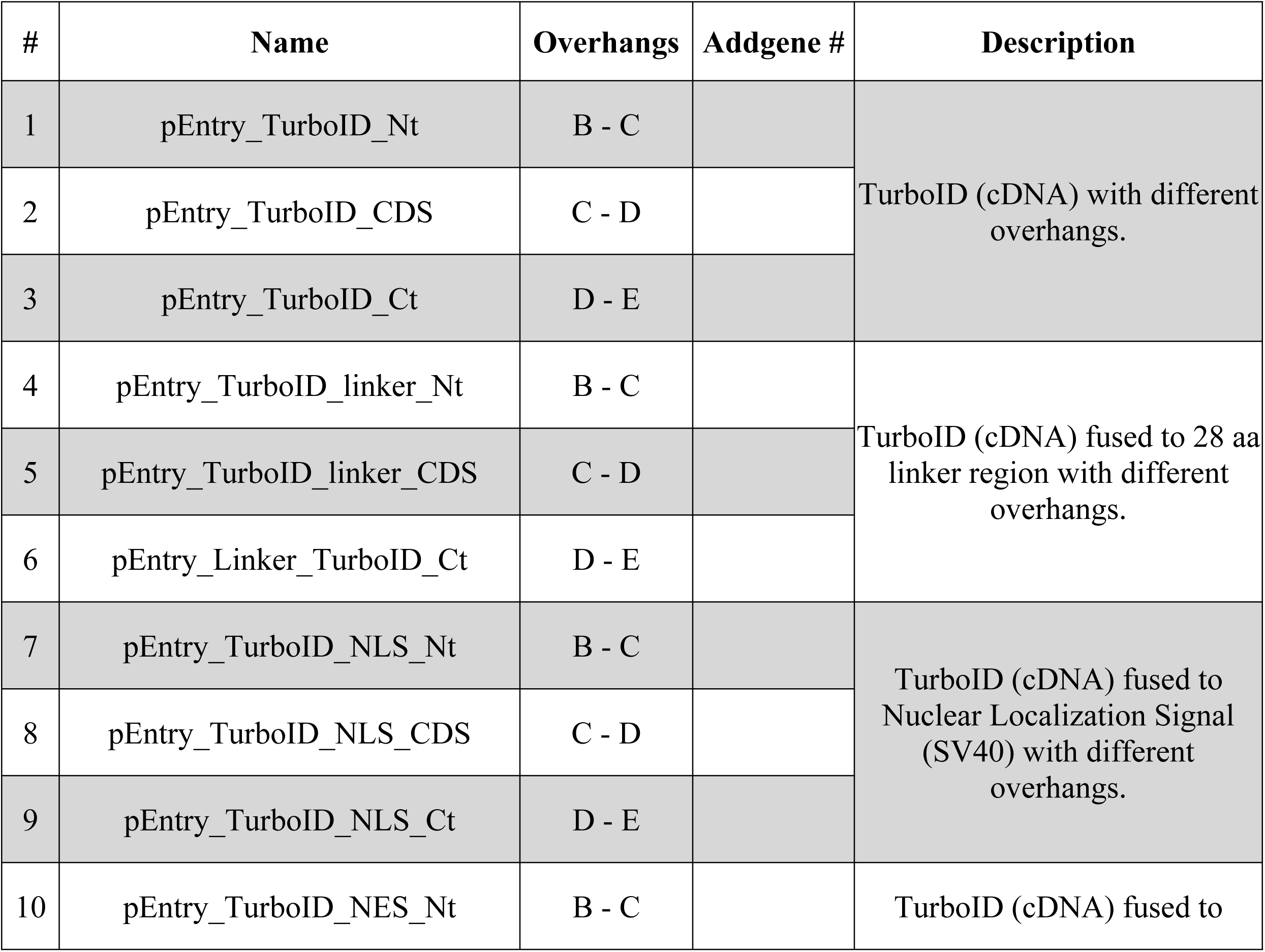
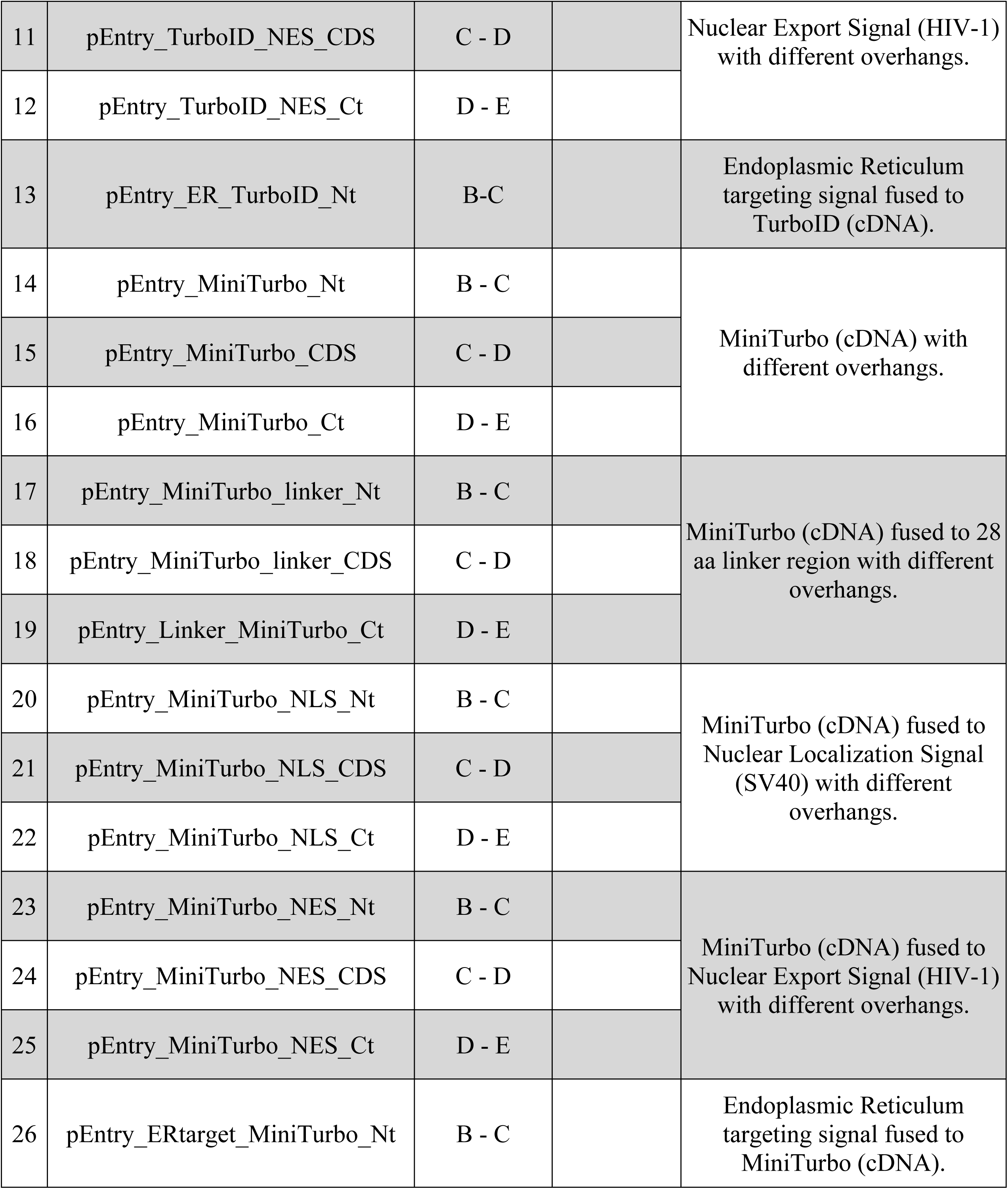
A plasmid collection for proximity labelling. Plasmid names include the Green Gate terms ‘Nt’, ‘CDS’ or ‘Ct’ for different overhang pairs generated after plasmid digestion with *Bsa*I. Linker refers to flexible 28 aa long sequence, NLS to Nuclear Localization Sequence (SV40), NES to Nuclear Export Signal (HIV-1) and ER to Endoplasmic Reticulum targeting signal. pEntry vectors designated as ‘Ct’ vectors (overhangs D-E) contain a stop codon at the end of the cassette. Vector maps are available at Addgene under the accession numbers provided.

The ability of biotin ligase to label an interactor is dependent on the proteins being proximal. In the case of a large protein, or a large protein complex, the labelling radius of a ligase fusion may not be sufficient to label interacting proteins [17]. The addition of flexible linker regions have previously been shown to increase the biotinylation range of a biotin ligase from its fused protein of interest [17]. Therefore, we included module constructs encoding each of the two ligases with an additional 28 aa long flexible linker region.

The inclusion of a biotin ligase-only control, whereby the biotin ligase is expressed without the protein of interest, is useful to interpret the potentially large list of peptides identified in a proximity labelling experiment [18]. This allows for the identification of proteins that may have been biotinylated because they are highly abundant in the cell or have some affinity to the biotin ligase protein itself. In order to identify physiologically relevant interaction partners this protein should be expressed at the same level as the biotin ligase-gene of interest and should also be localized to the same subcellular compartment. To allow for some change in ligase localization we generated biotin ligase constructs encoding the ligases fused to a nuclear localization signal (NLS) derived from simian virus 40 (SV40) T antigen [19], a nuclear export signal (NES) from human immunodeficiency virus 1 (HIV-1) protein Rev [20] and an N-terminal signalling peptide from sweet potato sporamin targeting the protein to the endoplasmic reticulum (ER) [21]. To facilitate community use, all plasmids have been deposited with the Addgene repository (see Table 1; Materials and Methods).

### TurboID fusion proteins are active in tobacco leaves and localize to different subcellular compartments

To ensure that the different localization signals do not interfere with biotin ligase activity and that they drive localization of the protein to the predicted cellular compartment we generated, using the entry modules described above, constructs expressing TurboID under the UBQ10 promoter (UBQ10p) fused to Green Fluorescent Protein (GFP) and encoding one of the localization signals (see Fig. 2A; Materials and Methods). These constructs were then transformed into tobacco leaf cells by *Agrobacterium*-infiltration. Using immuno-blotting we found that the expression of TurboID fusion proteins led to higher levels of biotinylated proteins in total protein extracts when compared to untransformed plants, likely as a consequence of biotin being naturally present in plant cells [6]. The signal for biotinylated proteins increased upon addition of extracellular biotin in to the leaf, consistent with previously described TurboID experiments in tobacco (Fig. 2B) [6].

**Fig 2.**
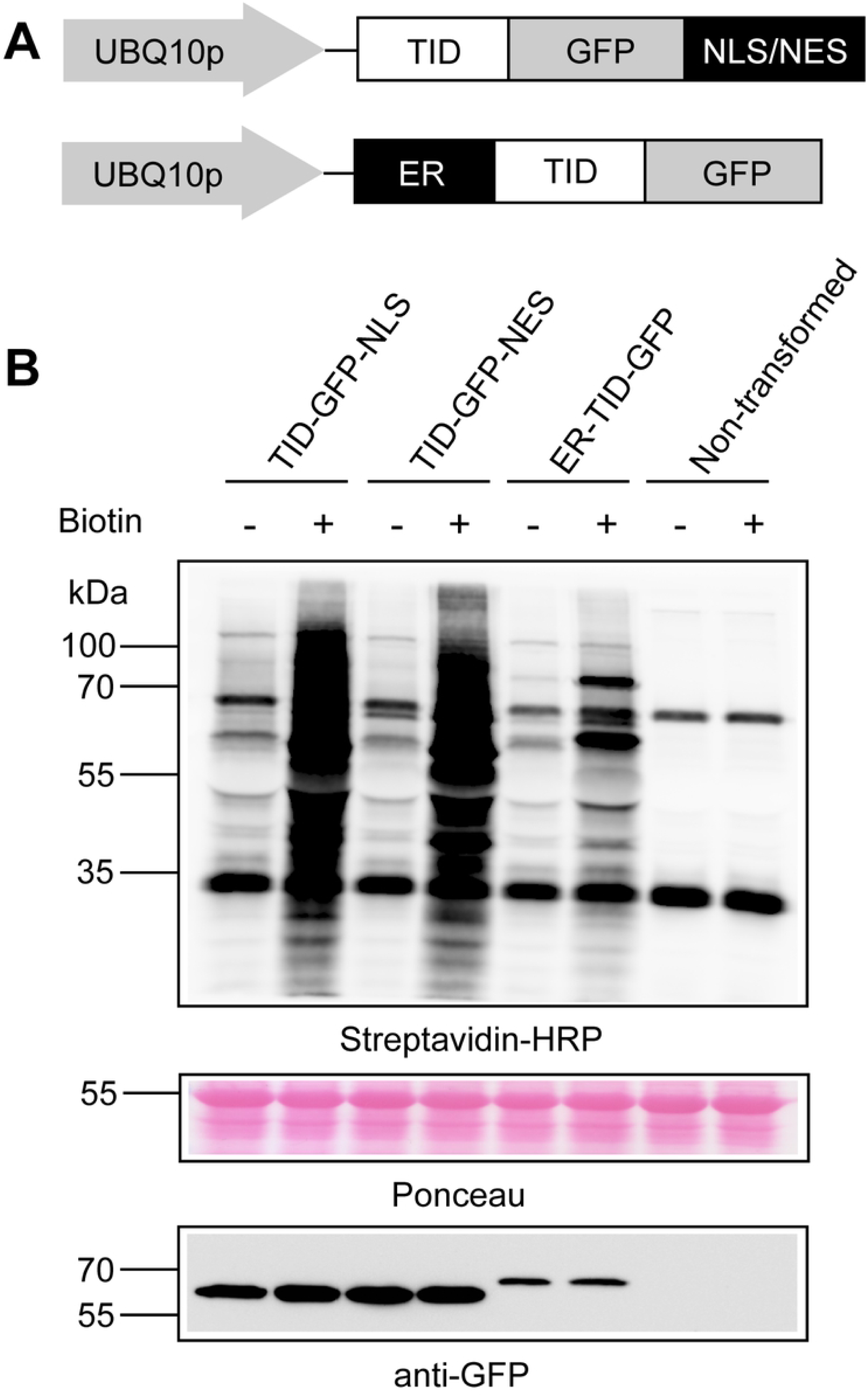
TurboID fusion proteins are active in tobacco leaves. **(A)** Schematic representation of the TurboID-GFP fusion proteins expressed in tobacco leaves. **(B)** Expression of the different TurboID-GFP fusion proteins in tobacco leaves increases the abundance of biotinylated proteins. Total protein extracts were prepared from non-trnasformed *N. benthamiana* leaves as well as from leaves transiently expressing the TurboID-GFP fusion proteins. TurboID fusion proteins were detected by immunoblotting using anti-GFP antibodies while biotinylated proteins were detected using streptavidin-HRP. TurboID-GFP-NLS/NES and ER-TurboID-GFP are expected to migrate at ∼64 kDa and 66 kDa, respectively. Experiment was conducted twice with similar results.

To observe the localization of the different TurboID-GFP fusion proteins we took sections of tobacco leaves transiently expressing the constructs and examined their fluorescence pattern using confocal microscopy. TurboID fused with GFP and the different localization signals gave fluorescent signals consistent with their targeted subcellular localization to the nucleus, cytoplasm and endoplasmic reticulum when compared with non-transformed leaves (Fig. 3). Taken together these results indicate that TurboID constructs generated with the library entry modules are active *in planta* and allow for the efficient targeting of TurboID – protein fusions to different subcellular compartments.

**Fig 3.**
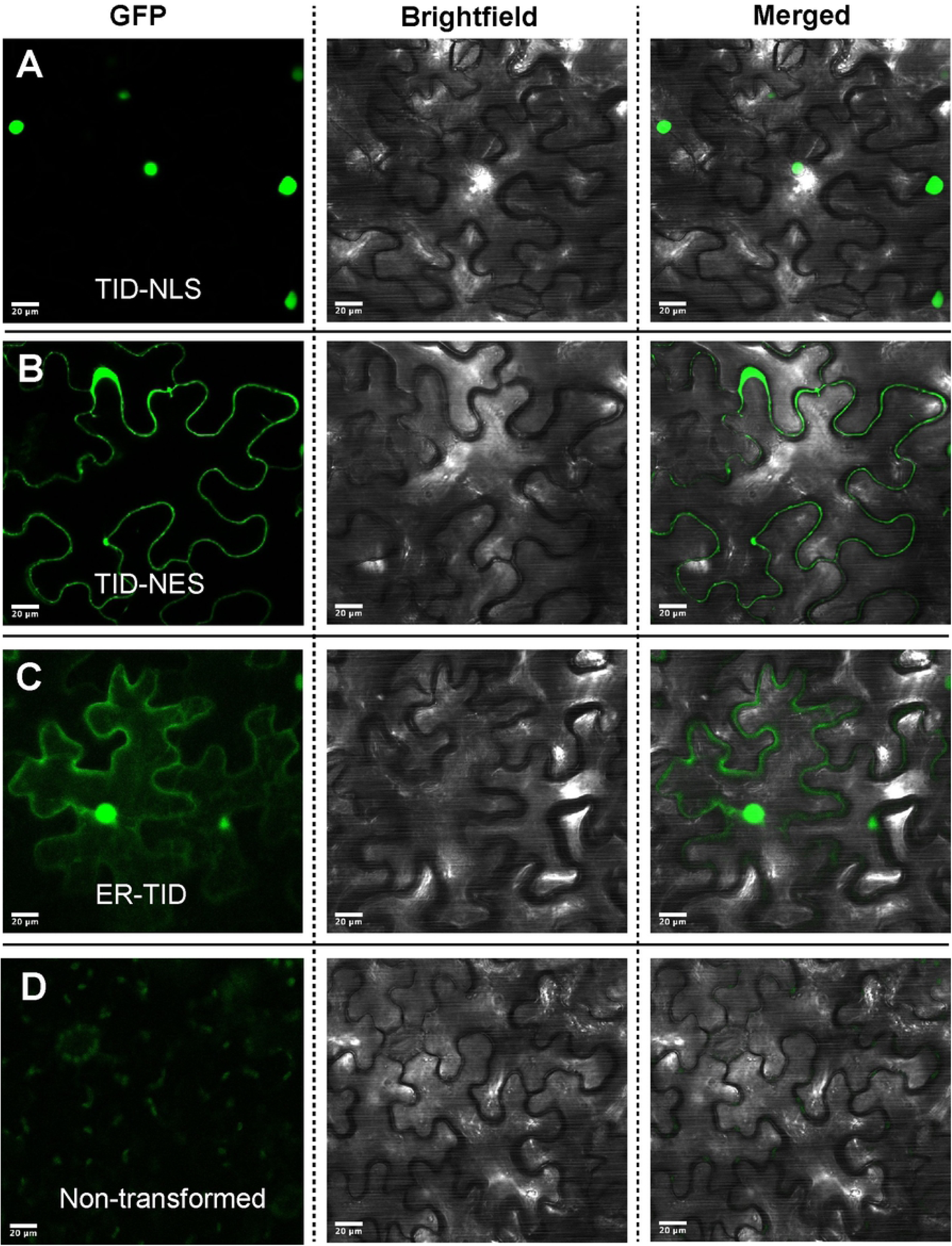
Subcellular localization of TurboID-GFP fusion proteins. Confocal microscopy images of abaxial epidermal cells from *N. benthamiana* leaves 3 d after infiltration with *A. tumefaciens* containing TurboID-GFP (TID) constructs with **(A)** a Nuclear-Localization Signals (NLS), **(B)** a Nuclear-Export Signal (NES), or **(C)** an Endoplasmic Reticulum targeting signal (ER). In **(D)**, abaxial epidermal cells from non-transformed leaves are shown. Scale bars: 20 μm.

### TurboID fusion proteins efficiently label interaction partners

We next tested if a protein-TurboID fusion construct generated with the entry modules resulted in the labelling of physiologically relevant interaction partners. To this end we performed a co-expression experiment in tobacco leaves using two *Arabidopsis* MADS domain transcription factors, *APETALA3* (*AP3*) and *PISTILLATA* (*PI*), which are known to physically interact with one another and function as an obligate heterodimer during flower development [22, 23]. Tobacco leaves co-expressing an AP3-GFP construct with a PI-TurboID fusion protein (PI–TurboID–3xMyc) (Fig. 4A) were infiltrated with either a mock or biotin-containing solution and tissue was collected after 3 hours. As AP3 and PI co-localize to the nucleus we generated nuclear protein extracts (see Materials and Methods). After immunoprecipitation of AP3-GFP using anti-GFP antibody coated beads a band at approximately 55 kDa could be observed corresponding to biotinylated AP3-GFP (Fig. 4B). This band was strongest in biotin-treated leaves but was also present in leaves not treated with exogenous biotin (Fig. 4B). This suggests that endogenous biotin in tobacco leaves is sufficient for some labelling of proximal proteins, albeit to a lesser extent. Another band, which migrated at the same size as PI-TurboID-3xMyc (∼65 kDa), was observed on the membrane (Fig. 4B). This is likely auto-biotinylated PI-TurboID-3xMyc which co-immunoprecipitated with AP3-GFP, consistent with previous reports showing biotin ligase auto-biotinylation [3, 24]. These results demonstrate that the entry modules generated for TurboID cloning can be used to construct a TurboID fusion protein that labels physiologically relevant interaction partners *in vivo*.

**Fig 4.**
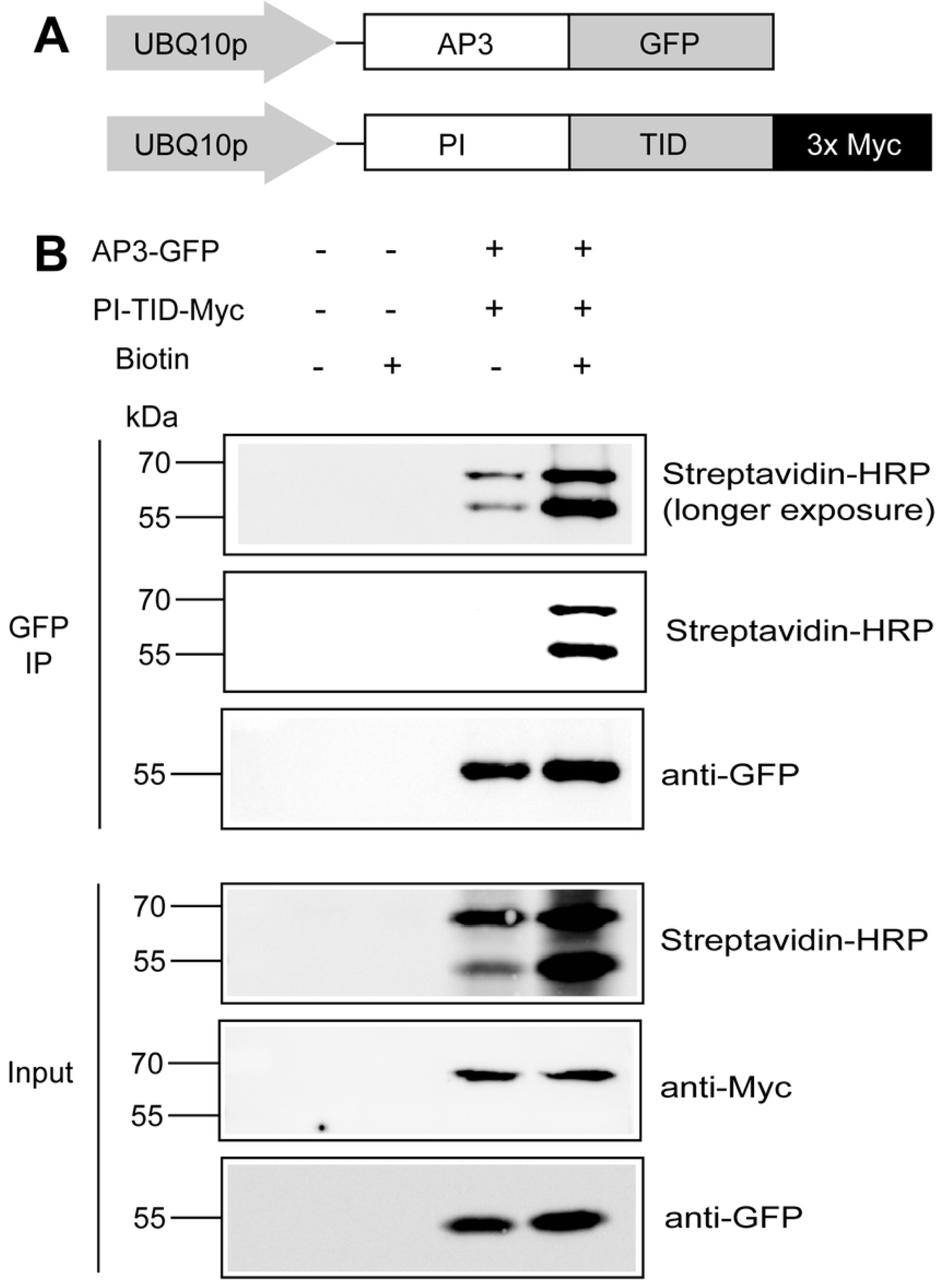
TurboID expression construct labels interaction partners of bait protein. **(A)** Schematic representation of the fusion proteins expressing the known interactors AP3 and PI fused to GFP and TurboID-3xMyc, respectively, expressed in tobacco leaves. **(B)** Biotinylation of AP3-GFP by PI-TurboID-Myc. Nuclear protein extracts were prepared from non-transformed *N. benthamiana* leaves or from leaves transiently expressing fusion the proteins. Immunoprecipitates were obtained using anti-GFP coated beads (ChromoTek). AP3-GFP, PI-TurboID-Myc and biotinylated proteins were detected using anti-GFP and anti-Myc antibodies and streptavidin-HRP, respectively. AP3-GFP is predicted to migrate at ∼55 kDa, PI-TurboID-Myc at ∼65 kDa. This experiment was conducted twice using different anti-GFP coated beads (ChromoTek GFP-Trap or Miltenyi μMACS Magnetic beads) with similar results. In the upper-most panel, a longer exposure (5 min) was chosen to better visualize faint bands appearing in the absence of exogenous biotin treatment in an extract from leaves expressing both the AP3-GFP and PI-TurboID-Myc fusion proteins.

## MATERIALS AND METHODS

### Plant growth conditions

*N. benthamiana* plants were grown on a medium consisting of compost, perlite and vermiculite in a ratio of 3:1:1. The plants were grown under constant illumination at 20°C after being incubated at 4°C in the dark for 5 d.

### Generation of biotin ligase cloning modules

All cloning procedures are described in the Supplementary Materials. Oligonucleotides used and plasmids generated are described in Table S1 and S2, respectively.

### Transient gene expression in *N. benthamiana*

*Agrobacterium tumefaciens* cells (C58 pGV2260 containing pSOUP) were transformed with plasmid DNA and plated on LB plates supplemented with appropriate antibiotics for 2-3 d at 28°C. A colony from this plate was used to inoculate LB medium supplemented with appropriate antibiotics and grown overnight at 28°C with shaking (∼225 rpm). *A. tumefaciens* cells were centrifuged at 4,000 *g* for 10 min and resuspended at an OD_600_ of 0.75 in Infiltration Medium (10 mM MES pH 5.6, 10 mM MgCl_2_, 150 μM acetosyringone). Approximately 4-5 week old *N. benthamiana* leaves were infiltrated with this solution using a blunt 1 mL syringe and leaf tissue was examined or stored in N_2_(l) after 3 d.

### Protein extraction and immunoblotting

To prepare total protein extracts from *N. benthamiana* leaves for immunoblotting, plant tissue was ground to a fine powder in N_2_(l). This powder was then resuspended in standard 2X SDS-loading buffer and boiled for 5 min (95°C). Samples were centrifuged and the supernatants were used for SDS-PAGE and immunoblotting. Antibodies were diluted in 1X phosphate-buffered saline plus Tween 20 (0.05%) with either 5% (w/v) milk powder (for anti-GFP and anti-Myc immunoblots) or 5% Bovine Serum Albumin (BSA) powder (streptavidin-HRP immunoblots). Primary antibodies used were: anti-GFP [1:5,000], Roche #1181446001; anti-Myc [1:1,000], #M5546, Sigma; streptavidin-HRP [0.2 μg/ml], Thermo Fisher Scientific #S911.

### Confocal imaging to visualize TurboID-GFP subcellular localization

In order to visualize GFP-tagged proteins expressed in tobacco leaves a *Zeiss* LSM 710 laser scanning confocal microscope was used. Tobacco leaf tissue was placed on a slide with H_2_O to prevent excessive drying and covered with a coverslip. Protein localization was visualized from the abaxial leaf side using a 488 nm excitation wavelength. Image processing was performed in FIJI (Image J) [25].

### *N. benthamiana* nuclear enrichment and AP3-GFP immunoprecipitation

Three days after *Agrobacterium*-infiltration of tobacco leaves (as described above) leaf tissue was infiltrated with a biotin solution (250 μM biotin, 10 mM MgCl_2_) or a mock solution (10 mM MgCl_2_) using a blunt 1 mL syringe. After 3 h of labelling leaves were frozen in N_2_(l) and ground to a fine powder. The powder was resuspended in 30 mL buffer M1 (10 mM sodium phosphate, pH 7, 0.1 M NaCl, 1 M 2-methyl 2,4-pentanediol, 10 mM β-mercaptoethanol and 1X Complete Protease Inhibitor Cocktail (CPIC, Sigma #1169749800). The solution was left at 4°C on a rotator for 10 min before being filtered twice through Miracloth (Merck Millipore) and spun at 1,000 *g* for 20 min at 4°C. The pellet was gently resuspended in 5 mL ice-cold M2 buffer (M1 buffer with additional 10 mM MgCl_2_ and 0.6% Triton X-100) before being centrifuged at 1,000 *g* for 10 min at 4°C. Pellet washes with M2 buffer were repeated 3 times. The pellet was then gently resuspended in 1 mL M3 buffer (10 mM sodium phosphate, pH 7, 0.1 M NaCl, 10 mM β-mercaptoethanol and 1X CPIC) and centrifuged at 1,000 *g* for 10 min at 4°C. The pellet was resuspended in 300 μL lysis buffer (150 mM NaCl, 1% Triton X-100, 50 mM Tris HCl pH 8.0, 0.1% SDS, 0.5% sodium deoxycholate) before being sonicated using a Diagenode Bioruptor Pico for 10 cycles of 30 s on/off at 4°C, and centrifuged at 12,000 x *g* for 5 min at 4°C. Protein concentration was determined using an Amido Black assay[26]. Protein extracts were diluted to 800 μL in IP buffer (150 mM NaCl, 1% Triton X-100, 50 mM Tris HCl pH 8.0). Anti-GFP coated beads were added (25 μL ChromoTek GFP-Trap or 50 μL MACS magnetic) and IP was carried out as per the manufacturers’ instructions. Proteins were eluted from the beads by addition of 2X SDS-loading buffer and incubation for 5 min at 95°C. The eluate was stored at -20°C until being analyzed by immunoblotting.

### Availability of the plasmid vectors

All constructs for proximity labelling have been deposited with Addgene under accession numbers XXX (see Table 1) and are available from there.

## Acknowledgements

We thank Dr. Brendan Davies for discussions and Dr. Emmanuelle Graciet for help with experiments. We also thank Drs. Chris Greene and Matthew Campbell for help with confocal microscopy. Funding for this work was provided by Science Foundation Ireland and the Environmental Protection Agency (grant 16/IA/4559 to F.W.).

